# Proteome-wide landscape of solubility limits in a bacterial cell

**DOI:** 10.1101/2021.10.05.463219

**Authors:** Ádám Györkei, Lejla Daruka, Dávid Balogh, Erika Őszi, Zoltán Magyar, Balázs Szappanos, Gergely Fekete, Mónika Fuxreiter, Péter Horváth, Csaba Pál, Bálint Kintses, Balázs Papp

**Affiliations:** HCEMM-BRC Metabolic Systems Biology Lab, Szeged, Hungary; Biological Research Centre, Institute of Biochemistry, Synthetic and Systems Biology Unit, Eötvös Loránd Research Network (ELKH), Szeged, Hungary; Doctoral School in Biology, Faculty of Science and Informatics, University of Szeged, Szeged, Hungary; Biological Research Centre, Institute of Plant Biology, Eötvös Loránd Research Network (ELKH), Szeged, Hungary; Department of Biomedical Sciences, University of Padova, Padova, Italy; Laboratory of Protein Dynamics, University of Debrecen, Debrecen, Hungary; Institute for Molecular Medicine Finland-FIMM, Helsinki Institute of Life Science-HiLIFE, University of Helsinki, Helsinki, Finland; HCEMM-BRC Translational Microbiology Research Group, Szeged, Hungary; University of Szeged, Department of Biochemistry and Molecular Biology, Szeged, Hungary

## Abstract

Proteins are prone to aggregate when they are expressed above their solubility limits, a phenomenon termed supersaturation. Aggregation may occur as proteins emerge from the ribosome or after they fold and accumulate in the cell, but the relative importance of these two routes remain poorly known. Here, we systematically probed the solubility limits of each *Escherichia coli* protein upon overexpression using an image-based screen coupled with machine learning. The analysis suggests that competition between folding and aggregation from the unfolded state governs the two aggregation routes. Remarkably, the majority (70%) of insoluble proteins have low supersaturation risks in their unfolded states and rather aggregate after folding. Furthermore, a substantial fraction (∼35%) of the proteome remain soluble at concentrations much higher than those found naturally, indicating a large margin of safety to tolerate gene expression changes. We show that high disorder content and low surface stickiness are major determinants of high solubility and are favored in abundant bacterial proteins. Overall, our proteome-wide study provides empirical insights into the molecular determinants of protein aggregation routes in a bacterial cell.

## Introduction

Maintaining solubility is a key requirement for proper functioning of proteins. In general, proteins are prone to misfold and aggregate at high concentrations due to the increased probability of forming intermolecular contacts that favor the aggregated state (Knowles *et al*, 2014). Thus, proteins become supersaturated and spontaneously aggregate when exceeding a critical concentration (Vecchi *et al*, 2020; Ciryam *et al*, 2013). Proteins with high supersaturation risks are those that are expressed at high concentrations with respect to their solubility limits (Ciryam *et al*, 2013). Such proteins are more likely to exceed their critical concentrations and aggregate under varying conditions. Indeed, recent proteomic studies established that supersaturated proteins are important sources of protein aggregates accumulating during ageing and protein homeostasis stresses (Vecchi *et al*, 2020; Sui *et al*, 2020).

Supersaturation may lead to aggregation of proteins from the nascent unfolded state (i.e. during or immediately after synthesis) or after initial folding and accumulation in the cell (Ciryam *et al*, 2013) (Figure 1). Proteins that are most at risk of supersaturation during their initial folding exceed their critical concentrations in their unfolded states as newly synthesized proteins. This route of aggregation may be avoided by quality control processes in the translational milieu, such as the ribosome-associated chaperon system (Young et al 2004). In sharp contrast, aggregation of already folded proteins is related to the inability of the cell to maintain proteins in their functional soluble states and may be initiated by local unfolding events or structural fluctuations, exposing hydrophobic surfaces (Chiti & Dobson, 2009). As such, this route of aggregation has relevance to various fields from protein deposition diseases (Chiti & Dobson, 2009) to the design of proteins with enhanced kinetic stability (Broom *et al*, 2015). Despite its fundamental importance, however, we have only a limited understanding of which proteins are prone to aggregate from the folded rather than from the unfolded state. Prior proteomic studies identified *in vivo* supersaturated proteins, but not the route by which they aggregate (Vecchi *et al*, 2020; Sui *et al*, 2020). Addressing this gap requires a proteome-wide study of the solubility limits in the unfolded and folded states.

**Figure 1.**
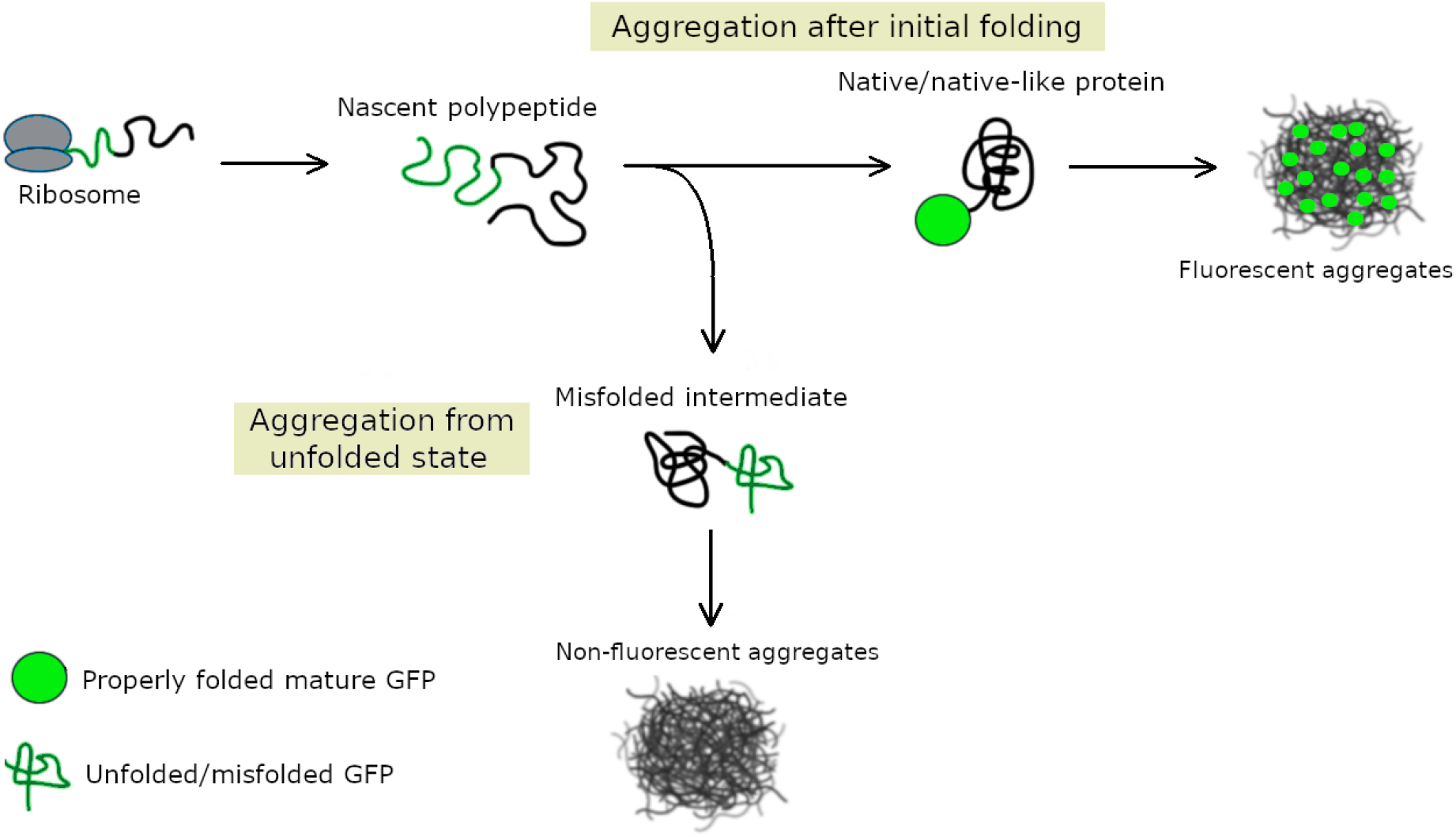
Two major aggregation routes in bacteria. Upon supersaturation, nascent polypeptides may aggregate from unfolded or partially folded intermediates through anomalous intermolecular interactions (lower pathway, black ball), without ever reaching their native structures (Groot *et al*, 2009). A fused GFP tag would show no fluorescence in this route because aggregation occurs before correct folding of the GFP moiety. Supersaturated proteins may also aggregate after reaching a native-like conformation as a result of local unfolding events (upper pathway) (Chiti et al, 2009). Because aggregation occurs after GFP chromophore formation, this route of aggregation would result in fluorescent aggregates (de Groot & Ventura, 2006).

Here we describe a high-throughput approach that systematically probes these two routes of aggregation in the model bacterium *Escherichia coli*. In this system, we overexpressed all proteins at similar very high levels thereby allowing comparison of the supersaturation potential across the whole proteome. The fate of overexpressed proteins was monitored by a GFP tag fused to the C-terminal of each protein. Such a GFP tag is a sensitive indicator of proper folding of the upstream protein (Waldo *et al*, 1999; Gregoire & Kwon, 2012; Zhang *et al*, 2005). Indeed, it has been widely used to improve the folding and solubility of a large variety of insoluble proteins in engineering applications (Waldo *et al*, 1999; Pédelacq *et al*, 2002). Importantly, while a lack of fluorescent signal indicates misfolding and aggregation of the upstream protein before the GFP chromophore is committed to form, a strong fluorescent signal indicates properly folded, soluble fused upstream proteins (Waldo *et al*, 1999). Whereas GFP has long been used to distinguish between these two groups of proteins, case studies revealed a third phenotypic group of GFP-fused proteins that form fluorescent aggregates (de Groot & Ventura, 2006; Rokney *et al*, 2009; Bakholdina *et al*, 2021). In particular, while rapid aggregation of an upstream peptide resulted in non-fluorescent aggregates, slow aggregation allowed correct folding of the GFP moiety and yielded fluorescent aggregates (de Groot & Ventura, 2006). These observations led us to hypothesize that the C-terminal GFP fusion informs on the route of aggregation, i.e. whether aggregation occurs right after synthesis or at a later stage, likely after folding. This question is important, as the genome-wide prevalence of these two routes of aggregation remains unknown.

Based on these considerations, we used high-content microscopy to evaluate the intensity and subcellular distribution of fluorescence signals in order to distinguish between overexpressed proteins that (i) remain highly soluble, (ii) aggregate rapidly from the newly synthesized unfolded state (aggregates without GFP signal) or (iii) aggregate later, potentially after initial folding (fluorescent aggregates). We then applied a data mining approach to systematically identify protein features that distinguish between these classes of proteins.

Our proteome-wide analysis shows that proteins in fluorescent aggregate have a low aggregation rate from the unfolded state, tend to fold rapidly and interact more frequently with the chaperone DnaK as compared to those in non-fluorescent aggregates. Together, these features indicate that these proteins likely aggregate after folding and there is a kinetic competition between aggregation and folding of the nascent polypeptide. Importantly, our screen suggests that aggregation from the folded state is the major route of insolubility of cytosolic proteins, challenging the traditional view that aggregates are dominantly formed by proteins that never reach their native structures (Baneyx & Mujacic, 2004). Furthermore, a substantial fraction (∼35%) of the proteome remain soluble at concentrations much higher than those found naturally, indicating that many bacterial proteins are not expressed at levels close to their solubility limits. High disorder content and low surface stickiness are major determinants of such high solubility and ultimately influence the functioning of abundant proteins. Overall, our work provides a global view of the solubility limits and aggregation routes in a bacterial cell.

## Results

### Proteome-wide measurement of solubility limits

We probed the solubility of 3,706 native *E. coli* proteins upon strong overexpression at the single-cell level using an image-based high-throughput screen (Figure 2A). To this end, we monitored the fate of overexpressed proteins using a GFP fused to the C-terminal of each protein (Kitagawa *et al*, 2005; Drew *et al*, 2001). As discussed above, the intensity and subcellular distribution of the fluorescence signal is indicative of the successful folding and solubility of the upstream protein. Specifically, we distinguish between three classes of proteins, indicating different routes of aggregation. First, a homogeneous GFP signal dispersed uniformly in the cells indicates a folded and soluble protein (Natan *et al*, 2018). Second, proteins without a GFP signal indicate aggregation that occurs before the GFP chromophore is committed to form (Natan *et al*, 2018; Waldo *et al*, 1999; de Groot & Ventura, 2006). These aggregates, referred to as ‘dark aggregates’, therefore likely correspond to aggregation from the unfolded state during or immediately after translation. Third, proteins with small fluorescent foci represent aggregation that occurs after the GFP chromophore is committed to form (de Groot & Ventura, 2006), potentially indicating aggregation after reaching the folded state. We refer to these aggregates as ‘fluorescent aggregates’.

**Figure 2.**
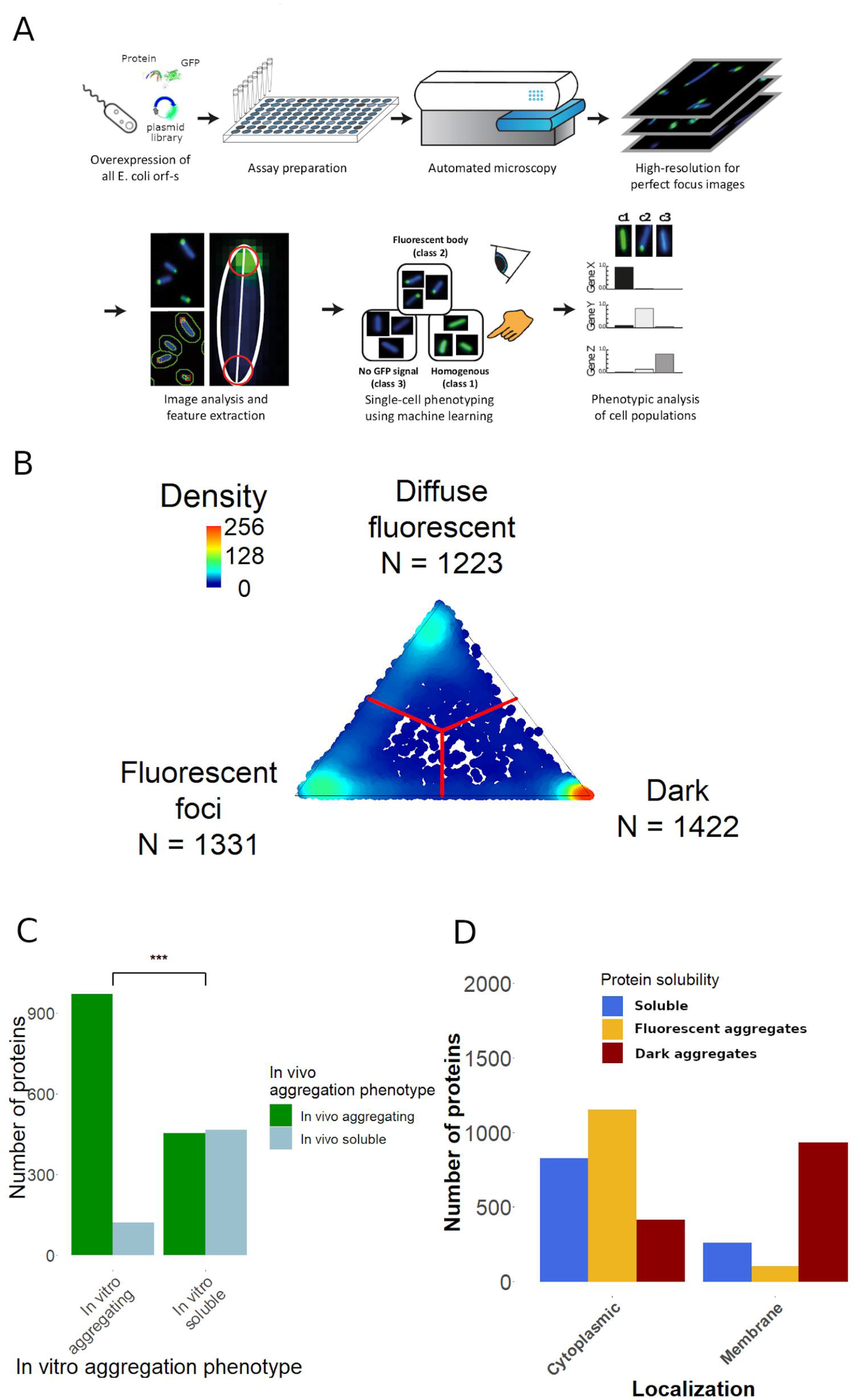
Experimental workflow and distribution of protein aggregation phenotypes. **A)** Workflow of high-throughput protein solubility measurement and classification. **B)** Distribution of cellular fluorescence phenotypes based on 1,000 classified cells for each overexpressed protein (represented by dots). The location of each dot is calculated from the fraction of cells showing each phenotype, with red lines representing the decision boundaries between aggregation categories assigned. The majority of proteins are located close to the vertices, demonstrating that cells typically show homogenous aggregation behavior. **C)** Comparison of in vivo and in vitro solubility phenotype of proteins. In vitro aggregating proteins show a strong overlap with proteins forming either dark or fluorescent aggregates in vivo (odds ratio=8.23, P < 10^-10^, Fisher’s exact test). **D)** Distribution of proteins according to their aggregation phenotypes and cellular localization. While 69% of membrane proteins form dark aggregates, only 17% of cytoplasmic proteins do so (P < 10^-10^, Fisher’s exact test).

After growing each overexpression strain in 96 well-plates, we applied a supervised machine-learning approach that automatically analyzes the images of approximately 1,000 cells for each overexpression strain (Figure 2A, Figure S1, Methods). Then, each overexpressed protein was classified into one of the above described three classes according to its predominant cellular phenotype: soluble, dark aggregate and fluorescent aggregate (Figure 2A, Table S1). We found that proteins of the *E. coli* proteome are distributed nearly equally among the three phenotypic categories (Figure 2B). Specifically, while 29.6% of the proteins were soluble despite the strong overexpression, 33.9% and 36.5% of the proteins formed fluorescent and dark aggregates, respectively.

To validate the accuracy of our high-throughput workflow, we took two complementary approaches. First, we assessed the overlap between the *in vivo* observed aggregation phenotypes and the *in vitro* measured solubility of the *E. coli* proteome, which was previously determined by expression using an *in vitro* reconstituted translation system (Niwa *et al*, 2009). Despite large differences in the *in vivo* and *in vitro* conditions, we found a strong overlap between the two datasets (odds ratio=8.23, P < 10^−10^, Figure 2C). In particular, 88.9% of the proteins that aggregate *in vitro* also show aggregation in our screen (Figure 2C). Because the *in vitro* dataset was generated without a GFP tag, the strong agreement between the two datasets indicates that the C-terminal GFP tag does not significantly influence the solubility of the fusion partners. Furthermore, when proteins aggregating *in vivo* were grouped on the basis of their fluorescence phenotypes (i.e. dark and fluorescent aggregates), the overlap between *in vitro* and *in vivo* solubility remained equally high across the two categories (Figure S1). Thus, the GFP tag is unlikely to bias towards one or the other class of aggregate. Second, to rule out that the expression levels of the proteins in the dark aggregate class are below the fluorescence detection limit, we measured the expression of a representative set of proteins by western blot analysis (see Methods; Table S2). We found that 95% of the tested proteins (i.e. 92 out of 97) from the dark aggregate class could be detected and showed comparable protein levels with those classified as fluorescent aggregates (Figure S2A and B). Crucially, 88.9% of these proteins are predominantly present in their aggregated forms inside the cells (Figure S2C). Finally, western blot analysis confirmed that 13 out of 14 proteins classified as soluble by the image-based screen are indeed present in the soluble cellular fraction (Figure S2C).

Taken together, these analyses indicate that our dataset is suitable to study the *in vivo* aggregation routes of overexpressed *E. coli* proteins on a proteomic scale.

### Cytosolic proteins have low risks of supersaturation from the unfolded state

We next compared the subcellular localizations of proteins under normal expression levels across the different aggregation classes. We report that the majority (69%) of membrane proteins form dark aggregates, while only a minute fraction (∼6%) of them form fluorescent aggregates (Figure 2D). This finding is in line with the notion that overexpressed membrane proteins are expected to aggregate from the unfolded state, before accessing the membrane where they would reach their native structure (Ciryam *et al*, 2013). This result also corroborates the high accuracy of our screen to detect aggregation that occurs from the unfolded state.

In sharp contrast to membrane proteins, only 17.2% of the cytoplasmic proteins form dark aggregates (Figure 2D), while 34.6% and 48.1% of them remain soluble and form fluorescent aggregates, respectively. The presence of fluorescent aggregates implies that at least a subpopulation of protein molecules remains soluble during or immediately after synthesis. As we overexpressed proteins well above their normal expression levels, these patterns indicate that most (70%) aggregation-prone cytosolic proteins must have a low risk of supersaturation from their unfolded states under physiological conditions.

### Molecular determinants of the two classes of aggregates

Based on prior case studies (de Groot & Ventura, 2006; Bakholdina *et al*, 2021), we hypothesize that fluorescent aggregates typically represent proteins that aggregate after initial folding. Owing to kinetic competition between aggregation and folding events, proteins that fold rapidly relative their rate of aggregation are expected to form aggregates after reaching their native state (de Groot *&* Ventura, 2006). A key prediction of this hypothesis is that proteins in fluorescent aggregates should have properties that prevent them from aggregation before their folding is completed. Analysis of the molecular properties that distinguish between proteins in dark versus fluorescent aggregates provides proteome-wide support for this hypothesis.

To systematically uncover the molecular determinants of the two classes of aggregates, we compiled a dataset of 115 protein features describing various physicochemical, structural and functional genomic properties for the vast majority of cytoplasmic *E. coli* proteins (see Table S3). Notably, predicted three-dimensional structures are available for 93% of cytoplasmic proteins in *E. coli* (Xu & Zhang, 2013). We focused on monomeric proteins only (N=1631) to avoid potential biases arising from oligomer interfaces (Pechmann *et al*, 2009). Next, we probed each feature for its ability to discriminate proteins that form dark and fluorescent aggregates from each other using logistic regression tests (Methods). The analysis revealed major differences between proteins in dark versus fluorescent aggregates that support the hypothesis (Figure 3, Table S4).

**Figure 3.**
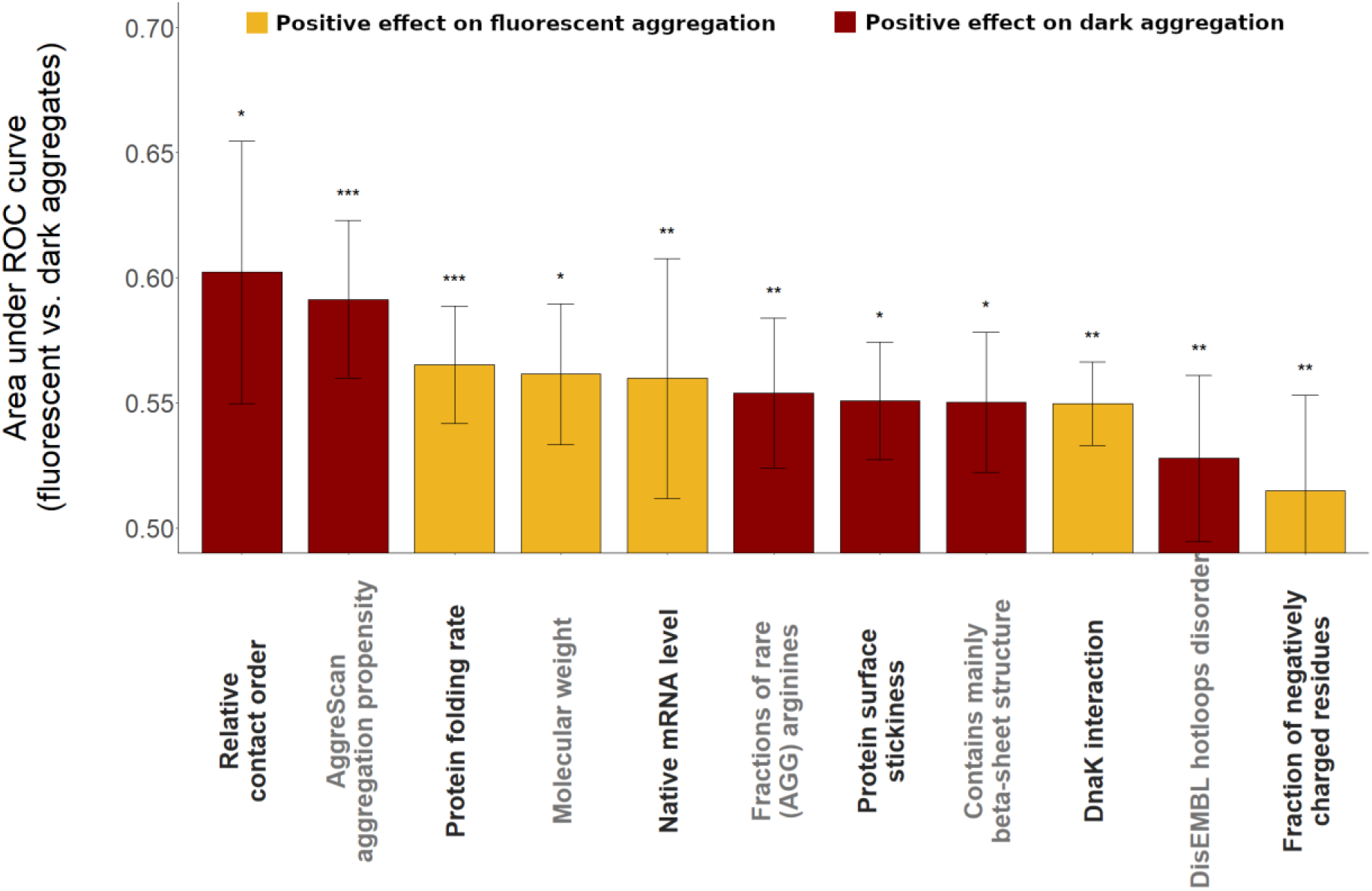
Major protein features distinguishing between the two routes of aggregation. Features discriminating between proteins that form dark versus fluorescent aggregates. The predictive ability of each feature was measured as the average area under the receiver operating characteristic (ROC) curve in a tenfold cross-validation procedure based on logistic regression analyses. All displayed protein features are statistically significantly predictive after adjustment for multiple testing using the false discovery rate method, ** corresponds to p_adj < 0.01, * to p_adj < 0.05 (logistic regression). Error bars show the 95% confidence interval for the AUC value of each feature.

First, proteins with more extensive aggregation hotspots (i.e. aggregation-prone sequence regions) are underrepresented among proteins in fluorescent aggregates compared to dark aggregates (Figure 4A, P = 5.01*10^−7^, logistic regression, based on the AggreScan (Conchillo-Solé *et al*, 2007) predictor, see Table S4 for other predictors). This is in line with previous results showing that the presence of aggregation hotspots increases the rate of aggregation from the unfolded state (de Groot *et al*, 2006; Chiti *et al*, 2003). As a further support, proteins in fluorescent aggregates show higher native mRNA levels than those in dark aggregates, suggesting that they have evolved a lower risk to aggregate from their unfolded state (Ciryam *et al*, 2013) (Figure 4B).

**Figure 4.**
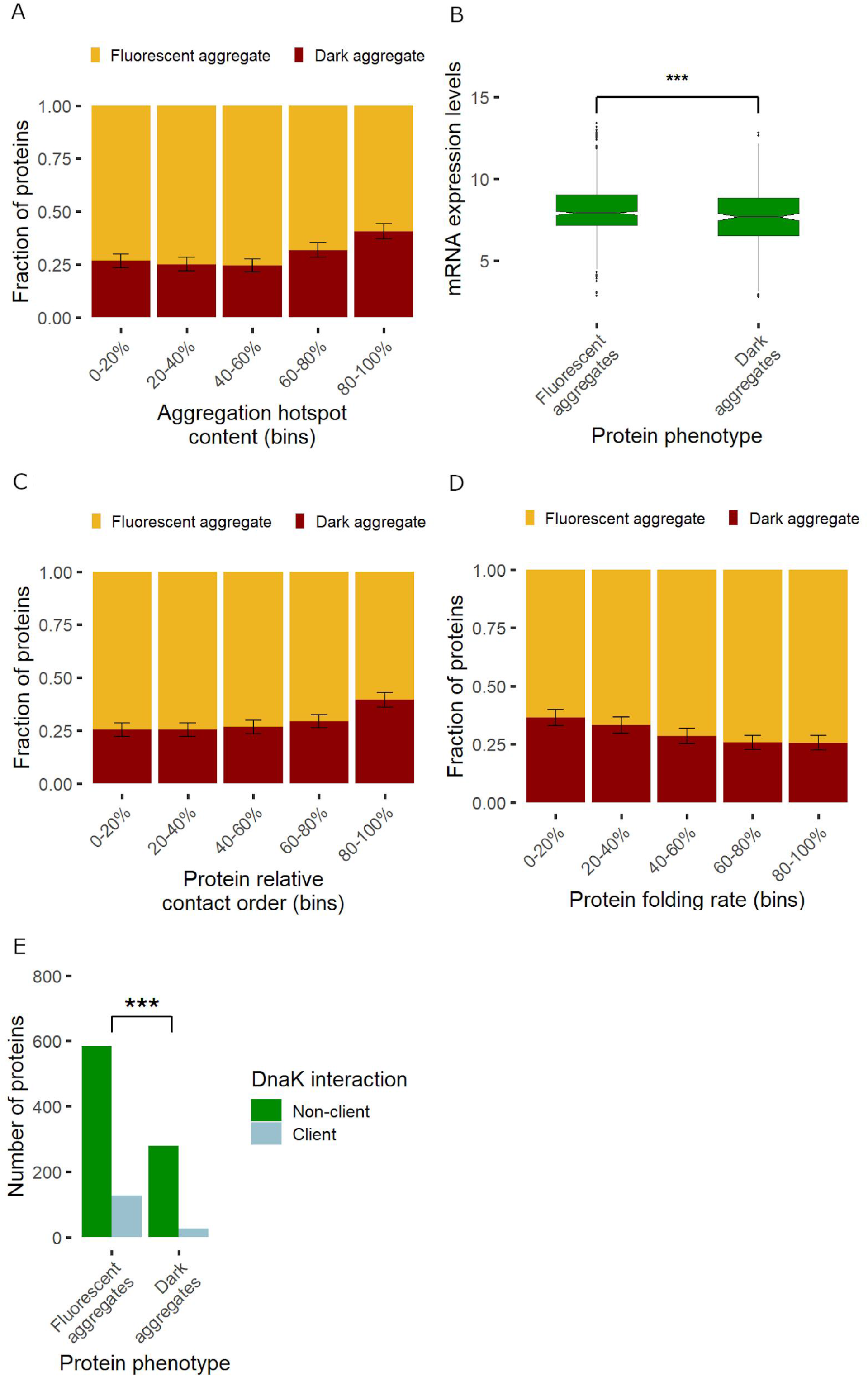
Aggregation and folding rates govern the two routes of aggregation. **A)** Proteins with more residues in aggregation hotspots, as estimated by AggreScan (Conchillo-Solé et al, 2007), are more likely to form dark than fluorescent aggregates (P = 5.01*10^-7^, logistic regression). **B)** Native mRNA expression levels of proteins in fluorescent aggregates are significantly higher than those in dark aggregate (P = 0.0008, Wilcoxon rank-sum test). **C**-**D)** Effects of protein contact order and folding rates on the class of aggregation. Note that a lower contact order and a higher folding rate (FOLD-RATE score) indicate easier folding. Dark aggregates are associated with a lower folding ability. **E)** Proteins in fluorescent aggregates are enriched in DnaK chaperone clients compared to those in dark aggregates (P = 4.94*10-5, Odds ratio = 2.43, Fisher’s exact test). Whiskers show standard errors and were calculated by bootstrap resampling.

Second, proteins in fluorescent aggregates tend to have higher folding rates, as indicated by two independent estimates (Figure 3 and 4C-D). We approximated folding rate by Fold-Rate (Gromiha *et al*, 2006), an ensemble predictor based on folding-correlated features derived from the primary sequence, and contact order (Plaxco *et al*, 1998), which measures sequence separation between contacting residues in the native state and is inversely correlated with folding rate (Methods). Both folding rate and contact order distinguishes between proteins in the two classes of aggregates (P = 0.0055 and P=0.004 by logistic regressions for folding rate and contact order, respectively, Figure 4C-D). Notably, contact order is the most predictive of the aggregation class out of the 115 tested protein features (Figure 3).

Third, proteins in fluorescent aggregates are more likely to be DnaK clients than those in dark aggregates (P = 4.95*10^−5,^ Fisher’s exact test, Figure 4E). DnaK is the major bacterial Hsp70 that interacts with hundreds of cytosolic proteins, most of which are assisted in their initial folding (Calloni *et al*, 2012). As DnaK promotes proper folding and prevents aggregation of client proteins (Balchin *et al*, 2020; Imamoglu *et al*, 2020), this result indicates that proteins in fluorescent aggregates are prevented from aggregation in their unfolded state.

Taken together, these findings are consistent with the notion that rapid folding and slow aggregation of a protein permits its folding before aggregation takes place (de Groot & Ventura, 2006). These results further suggest that fluorescent aggregates are typically formed by proteins that become supersaturated in their folded rather than in their unfolded state. Given that 70% of insoluble proteins display fluorescent aggregates in our screen (Figure 2D), we conclude that aggregation from the folded state is the predominant risk of supersaturation for cytosolic proteins in *E. coli*.

### Determinants of high solubility

Our proteome-wide screen revealed a subset of cytosolic proteins that remain soluble even when expressed well above their normal levels. Such highly soluble proteins are unlikely to be expressed at concentrations close to their solubility limits, hence are not supersaturated under physiological conditions. We next sought to identify the molecular properties that distinguish highly soluble proteins from both groups of aggregating proteins (Methods). Consistent with earlier works, soluble proteins show high mRNA expression levels (Tartaglia *et al*, 2009), high content of negatively charged residues (Niwa *et al*, 2009; Kramer *et al*, 2012), low content of hydrophobic amino acids (Giasson *et al*, 2000; Schwartz *et al*, 2001) and small size and surface area (Niwa *et al*, 2009) (Figure S3). More remarkably, our analysis identifies low surface stickiness and high protein disorder content as major determinants of solubility.

Proteins differ in their propensity to form non-specific interactions with other macromolecules depending on the ‘stickiness’ of their surfaces (Levy *et al*, 2012). While low stickiness has been linked to avoidance of non-functional interactions (Levy *et al*, 2012), its role in avoiding aggregation has remained unclear. We find that soluble proteins show a markedly lower surface stickiness score than either class of aggregates (P < 10^−10^ in both cases, Wilcoxon Rank Sum test, Fig 5A, see Methods). Notably, while highly abundant proteins tend to have less sticky surfaces (Levy *et al*, 2012), the association between high solubility and low stickiness remains when controlling for abundance (P = 2.38*10-7 and P = 0.00946 for fluorescent and dark aggregates, respectively, logistic regressions).

**Figure 5.**
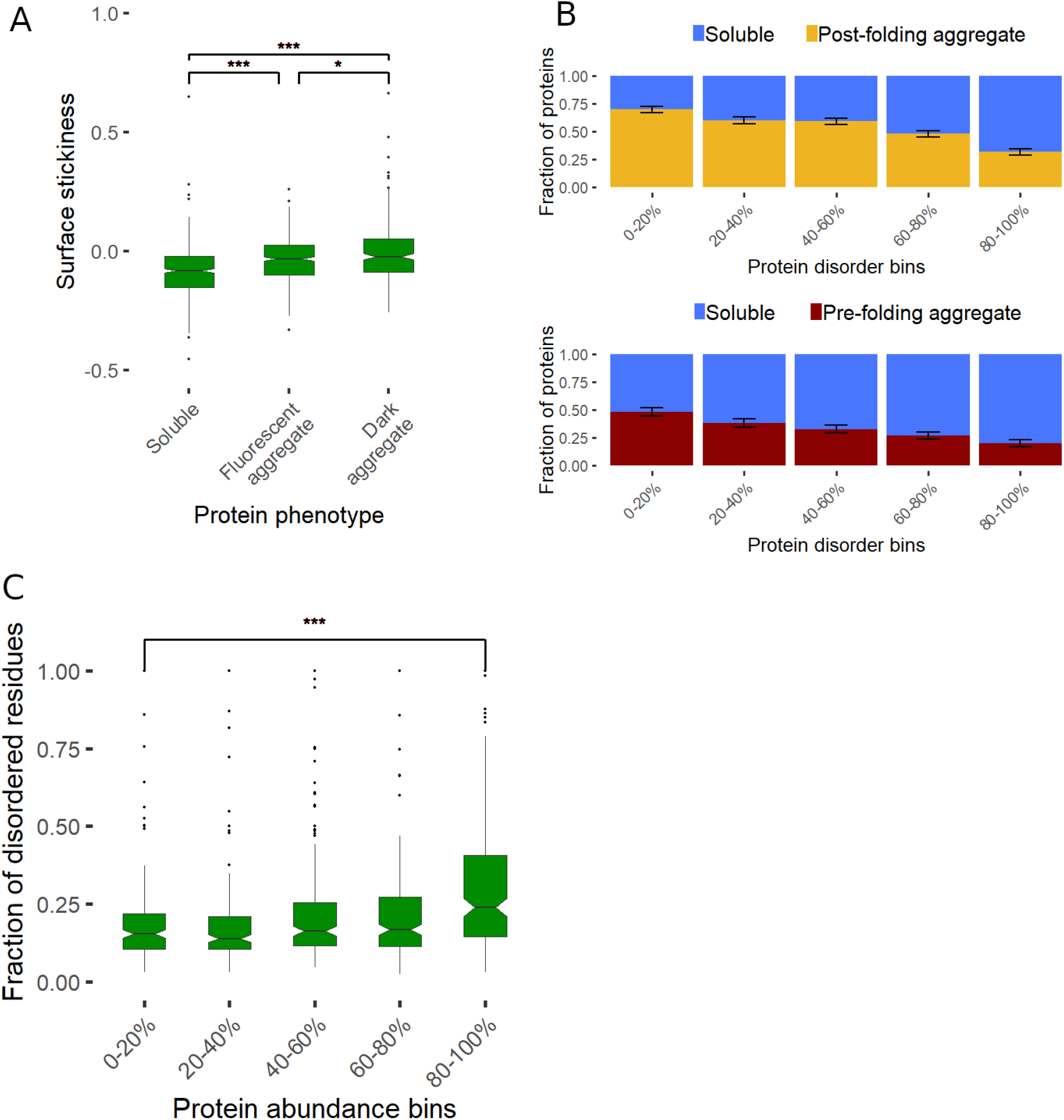
Protein stickiness and disorder content shape solubility. **(A)** We use the surface stickiness score to measure promiscuous interaction propensity (Levy et al, 2012). Both fluorescent and dark aggregates show higher surface stickiness than soluble proteins (P < 10^-10^ and P < 10^-10^ respectively, Wilcoxon Rank Sum test). **B)** Fractions of aggregating proteins as a function of disorder content (binned data), calculated using PONDR VSL2B. Upper panel shows the fraction of proteins in fluorescent aggregates among those that are either soluble or in fluorescent aggregates, while the lower panel shows the fraction of proteins in dark aggregates among those that are either soluble or in dark aggregates. **C)** Disorder content as a function of native protein abundance in E. coli based on (Schmidt et al, 2016). The most abundant 20% of E. coli proteins have a significantly higher disorder content than those in the least abundant 20% bin (P = 4.18*10^-11^, Wilcoxon rank sum test). Disorder content was calculated using PONDR VSL2B, but similar results are obtained with other predictors (see Tables S5 and S9)

Proteins often contain intrinsically disordered regions that lack a unique structure (Tompa, 2002). We find that the proportion of disordered residues (disorder content) of a protein is strongly associated with its solubility upon overexpression (Figure 5B). To examine the impact of disordered regions on this association, we next identified segments of contiguously disordered amino acids (≥10 residues). We find that soluble proteins contain both more such disordered segments per length and a higher proportion of disordered residues outside these segments than those forming either class of aggregates (Table S5). These results are robust to using different predictions of disorder (Table S5-S6) and are not simply a byproduct of other protein features associated with solubility (Table S7). Importantly, our analyses show that both disorder content and surface stickiness have independent effects on solubility (Table S8).

If disorder content and protein stickiness shape protein solubility at native expression levels as well then highly abundant proteins should exhibit high disorder content and low surface stickiness as evolutionary adaptations to enhance their solubility. Indeed, it has been reported that highly expressed proteins in *E. coli* exhibit low surface stickiness (Levy *et al*, 2012) and elevated disorder content (Paliy *et al*, 2008), albeit these patterns have not been linked to protein solubility. By analyzing the same set of cytosolic proteins for which aggregation phenotype was measured and using state-of-the-art proteomics data on native protein levels (Schmidt *et al*, 2016), we confirm these correlations (Table S9). Notably, the correlation between protein abundance and disorder content is mainly caused by elevated disorder in the most abundant group of proteins (Figure 5C), which is consistent with the notion that such proteins are highly optimized to avoid aggregation.

Collectively, these results suggest that elevated structural disorder and low surface stickiness enhance protein solubility in the bacterial cell and play important roles under physiological conditions.

## Discussion

Here we systematically probed the solubility limits and aggregation routes of *E. coli* proteins by overexpressing them and monitoring their fate using an image-based *in vivo* assay. We have identified on a proteome-wide scale those proteins that aggregate before reaching their folded state and those that likely aggregate after initial folding.

We found marked differences in the properties of proteins that become insoluble via the two routes of aggregation. Proteins aggregating after initial folding exhibit faster folding kinetics and a lower content of aggregation hotspots compared to proteins that aggregate from the unfolded state. We find that interaction with the DnaK chaperone also helps to reach the folded state in the translation milieu before aggregation. These observations provide empirical evidence that supersaturation may occur at different stages of a protein’s lifetime (Ciryam *et al*, 2013) and a relatively slow aggregation and rapid folding of a protein permit its folding before aggregation occurs. Overall, these findings also provide proteome-wide support for the notion that kinetic competition between proper folding and aggregation from the unfolded state is one of the major forces that govern the route of aggregation (de Groot & Ventura, 2006).

Our proteome-wide study revealed that the majority (∼70%) of the aggregation-prone *E. coli* proteins in the cytoplasm are robust against aggregation during or immediately after synthesis but rather aggregate after initial folding. This finding challenges the traditional view that overexpression-induced aggregates are mainly formed by proteins that never reach their native states (Baneyx & Mujacic, 2004). Because we monitored aggregation at expression levels much higher than those found naturally, this result also indicates that most proteins are unlikely to be supersaturated in their unfolded states under natural conditions. Our findings suggest that both the inherent properties of protein sequences (rapid folding and shortage of aggregation hotspots) and interaction with the chaperone system contributes to high solubility during initial folding. As a consequence, supersaturation in the folded state emerges as the predominant risk of aggregation in bacteria.

Our results suggest that non-specific protein-protein interactions play an important role in shaping the solubility limits of proteins. It has been shown that natively abundant proteins have evolved an especially low surface stickiness to avoid non-specific interactions and hence interference with other proteins (Levy *et al*, 2012). Our work goes further and demonstrates that proteins with low surface stickiness also tend to avoid aggregation. Based on these observations, we propose that avoidance of non-specific interactions is an important mechanism to reduce the aggregation propensity of highly expressed proteins.

Our study provides new insights into the role of disorder content in protein aggregation in two ways. First, we show that disorder content is a major driver of solubility differences between *E. coli* proteins *in vivo* independently of the effect of surface stickiness. Importantly, high disorder content is associated with solubility regardless of the overall charge and hydrophobicity of the proteins (Table S8), suggesting that it is the flexibility conferred by disordered regions what matters for enhanced solubility. This notion is consistent with prior works on the role of flexible structural elements in protein solubilization (Simone *et al*, 2012; Santner *et al*, 2014). Specifically, fusion of disordered segments to insoluble proteins have been shown to aid protein folding and solubilization by providing favorable surface area and by acting as “entropic bristles” (Santner *et al*, 2014). Our observation that soluble proteins also contain elevated frequencies of long disordered segments further supports the hypothesis that enhanced solubility is driven by entropic effects. Second, our work sheds new light on how natural selection shapes the disorder content of proteins. It has been noted earlier that highly expressed proteins display elevated disorder content in *E. coli*, however, the underlying selective constraints have remained puzzling (Paliy *et al*, 2008). Our results indicate that disorder content has evolved partly to avoid aggregation of highly abundant proteins under physiological conditions in *E. coli*. Thus, we propose that both low surface stickiness and elevated disorder content contribute to the high solubility of abundant proteins in bacteria (Tartaglia *et al*, 2009).

Our screen shows that about one third of cytosolic proteins in *E. coli* remain substantially soluble even when strongly overexpressed. This finding implies that these proteins are unlikely to be supersaturated under physiologically relevant conditions. This conclusion is broadly consistent with a prior proteomic study showing that about one quarter of proteins are expressed below their critical concentrations in *C. elegans* under non-stressed conditions (Vecchi *et al*, 2020). Importantly, our finding also has relevance to the ‘life on the edge’ hypothesis, which posits that proteins have evolved to be sufficiently soluble to allow their expression at the levels needed for their biological roles, but have almost no margin of safety to tolerate elevated concentrations (Tartaglia *et al*, 2007; Vecchi *et al*, 2020). Contrary to this hypothesis, the critical concentrations of soluble proteins identified in our screen appear to be much higher than those required to maintain their solubility under normal conditions. Thus, avoidance of supersaturation may not be the only evolutionary force affecting protein solubility for a subset of cytoplasmic proteins. We speculate that extremely high solubility might have evolved indirectly as a by-product of selection on other protein features that also influence solubility. For example, low protein surface stickiness might have primarily evolved to minimize unwanted protein interactions (Levy *et al*, 2012), but also enhances solubility as a by-product. Clearly, further studies are needed to decipher the evolution of extremely high critical concentrations.

Finally, the results presented in this study have biotechnological implications. Proteins that are prone to aggregate after reaching their folded states might form so-called non-classical inclusion bodies that contain folded proteins (Ventura & Villaverde, 2006) and may even show biological activities (García-Fruitós *et al*, 2005; Worrall & Goss, 1989). Such aggregates are of biotechnological importance as they offer efficient ways to recover active proteins and can be exploited as functional materials (Singhvi *et al*, 2019; García-Fruitós *et al*, 2012). From a methodological standpoint, our high-content microscopy based approach may serve as an efficient and scalable tool to screen for recombinant proteins that form aggregates after initial folding and hence may potentially contain folded proteins. An unresolved issue is the extent to which fluorescent aggregates identified in our screen contain catalytically active enzymes. Future efforts should address this issue by combining automated high-throughput inclusion body purification systems (Jäger *et al*, 2020) with large-scale functional profiling of enzymes (e.g. phosphatases) (Huang *et al*, 2015).

## Materials and methods

### Image-based high-throughput screen of protein aggregation

We used the C-terminal GFP fusion version of the *E. coli* K-12 Open Reading Frame Archive library (Kitagawa *et al*, 2005) to probe the aggregation pathways of 3,706 native *E. coli* proteins in an image-based screen. Part of this screen was previously published and included a set of 611 *E. coli* proteins that form homomers (Natan *et al*, 2018). In this previous work, we distinguished only two aggregation phenotypes (homogeneous GFP signal throughout the cells indicating a folded and soluble protein), and proteins without a GFP signal indicating aggregation before reaching the native conformations (that is, a “dark” aggregate). Here, we substantially extended this earlier dataset in two ways. First, here we also included homomers that show fluorescent foci and therefore represent inclusion bodies with properly folded C-terminally fused GFP (that is, a “fluorescent” aggregates). Second, we carried out the screen for the rest of the *E. coli* K-12 proteome. The applied protocol was as follows.

#### Cell preparation

The C-terminal GFP fusion version of the E. coli K-12 Open Reading Frame Archive library (Kitagawa *et al*, 2005) was grown in the original host strain *E. coli* K-12 AG1 in 96-well plates (growth conditions: 37°C, 280 rpm, LB medium supplemented with 20 µg/ml chloramphenicol as a selection marker of ASKA plasmids). We emphasize that this version of the GFP molecule is optimized to be highly expressed and show high fluorescence intensity in *E. coli* growing at 37°C (Kitagawa *et al*, 2005). Following overnight growth, expression was induced for 2 hours by 0.1 mM IPTG in the fully-grown culture at 37°C. From the induced cultures 0.2 μL were carried over using a pin tool replicator into black CellCarrier-96 plates (PerkinElmer). In this plate, each well had been supplemented with 100 μl of 5 μg/mL 4,6-diamidino-2-phenylindole (DAPI) in mineral salts minimal medium (MS-minimal) without any carbon source. Prior to the microscopic analysis, cells were centrifuged down to the bottom of the 96 well plates.

#### Imaging

Microscopy was done using a PerkinElmer Operetta microscope. Four sites were acquired per well. Laser-based autofocus was performed at each imaging position. Images of two channels (DAPI and GFP) were collected using a 60x high-NA objective to visualize the cell and the aggregation states of the proteins, respectively. At every site and every fluorescent channel 5 images were taken at different z positions with 0.5 μm shifts. These images were used for a perfect focus algorithm. Cellular properties of about 1000 cells of each expressing strain were extracted from the images, including the localization of the GFP signal within the cell.

#### Image Analysis

Images were pre-processed using the CIDRE algorithm (Goodsell & Olson, 2000) to remove uneven illumination. A perfect focus algorithm was developed to locally select the best z image plane and create an image that contains the highest contrast cells. To identify cells and extract their properties, the CellProfiler program (Carpenter *et al*, 2006) was used with custom modifications. First, image intensities were rescaled. Then, cells were identified on the DAPI signal using Otsu adaptive threshold and a Watershed algorithm to split touching cells. Cellular features such as intensity, texture, and morphology were extracted.

#### Phenotypic Classification using Machine Learning

Supervised classification of cells into predefined groups was performed using the Advanced Cell Classifier software (Piccinini *et al*, 2017). The cellular phenotypes were (i) no GFP signal (fluorescence level equals to that of the negative control without GFP) (ii) homogenous GFP signal (cells show equally distributed GFP signal throughout the whole cell) (iii) concentrated GFP signal in either one or both poles of the cell. Cells that did not fit into these three categories were discarded. For the automated decision, an artificial neural network method was used based on the Weka software (Hall *et al*, 2009).

Based on this cell classification, the proteins were assigned to one of the three classes, depending on which phenotype was predominant in the cell population. We considered a protein as “Dark” if the most populous category of cells showed no fluorescence. If the predominant cellular phenotype was a concentrated fluorescent spot at the cell pole, proteins were classified as “Fluorescent foci”, and if the majority of cells showed diffuse green fluorescence, proteins were classified as “Diffuse fluorescent”.

#### Protein expression analysis

We carried out western blot analyses for a representative set of protein overexpressions from the three groups of aggregation phenotypes with a special focus on the expression levels of the ‘dark aggregate’ group (i.e. those not showing a fluorescence signal). In brief, the C-terminal GFP fusion version of the *E. coli* K-12 Open Reading Frame Archive library (ASKA), members were grown overnight in 1 ml LB medium supplemented with 20 µM chloramphenicol at 37°C (Kitagawa *et al*, 2005). Following overnight growth, expression was induced for 2 hours by 0.1 mM IPTG in the fully-grown culture at 37°C. Following expression, cells were harvested by centrifugation (∼13,000g) and the pellets were resuspended in 250 µl 2xSDS-sample buffer. After boiling the samples for 5 minutes, 5 μl were separated on 10% SDS-polyacrylamide gel (PAGE). Gels were either stained with Coomassie Brilliant Blue (CBB) for justifying equal loading or transferred onto PVDF membranes (Amersham, GE Healthcare Lifescience) proceeding further for western blotting. Next, membranes were blocked in 5% (w/v) milk powder-0.05% (v/v) Tween20 in TBS (25mM Tris-Cl, pH 8.0, 150 mM NaCl) buffer (TBST) for an hour at room temperature (RT). Next, the membranes were incubated with 5% (w/v) milk powder-TBST including anti-GFP (Chromotek) as primary antibody (diluted to1:1000) and agitated overnight at 4°C. After washing three times with TBST buffer to remove the excess of unbound primary antibody, membranes were incubated with appropriate secondary antibody (Sigma-Aldrich) diluted in 2.5% (w/v) milk-powder-TBST buffer (1:10000) for an hour on RT. After washing the membranes three times in TBST buffer, signals were developed by a standard chemiluminescent western blot detection method (Thermo Scientific).

#### In vivo solubility analysis

Next, we tested the aggregation propensity of a representative set of protein over-expressions to confirm the predicted aggregation phenotypes coming from the microscopic image analyses. In brief, cells were grown as described above. After the centrifugation step, cell pellets were lysed by resuspending them in 200 μl BugBuster (Sigma) reagent at room temperature. Cell suspensions were incubated on a shaking platform at a slow setting for 20 min at room temperature. The solutions were centrifuged at 16,000 × g for 20 min at 4°C. 200 μl soluble fractions were removed and 50 μl 5xSDS-sample buffer was added. The pellets were resuspended in 250 µl 1xSDS-sample buffer. After boiling the samples for 5 minutes, 5 μl were separated on 10% SDS-polyacrylamide gel (PAGE). Western blot was carried out as described above. Signals were converted into black and white images and then quantification of the western blot bands (degradation products were not counted) was carried out by Image Studio Light. Band area was then corrected by eliminating the background value.

### Bioinformatics analysis

Each protein was assigned a cellular localization according to StepDB (Orfanoudaki & Economou, 2014), oligomerization data was retrieved from EcoCyc (Keseler *et al*, 2017). Only cytoplasmic monomers were used in further calculations, resulting in 1631 proteins. We collected 115 features for each protein, including mRNA level and protein abundance, presence of protein-protein interactions, disorder content, physico-chemical and functional properties from various sources (see Table S4 for details). Protein structures were retrieved from the Zhang Lab webpage (https://zlab.bio/) (Xu & Zhang, 2013), which were subsequently used in protein stickiness and surface area calculation as well as determining amino acid contents on the protein surface and folding calculations. Protein structure classification was retrieved from the CATH database (Sillitoe *et al*, 2019) and secondary structure superfamilies and families were used. Protein disorder data was retrieved from the MobiDB database (Piovesan & Tosatto, 2018) and calculated with several different predictors: DisEMBL (Linding *et al*, 2004), PONDR VSL2B (Peng *et al*, 2006), iUpred (Dosztányi *et al*, 2005) and Espritz (Walsh *et al*, 2012) with default parameters. The number of disordered amino acids was normalized to the protein length in all cases.

All 1631 proteins were used in subsequent calculations, with the notable exceptions of protein folding rate and relative contact order. These parameters can only be reliably calculated for single domain proteins, thus others were excluded, leaving 661 proteins. Area under receiver operating characteristic curve for logistic regression models was calculated with the pROC R package, with 10-fold cross validation. Enzyme Commission numbers were retrieved from the UniProt database.

All calculations were performed in R version 3.5.0, 2018-04-23 (R Core Team, 2008) in Rstudio version 1.1.447, figures were created in R base and ggplot2 version 2.2.1.

## Acknowledgements

We thank Péter Tompa for his valuable comments on an earlier version of the manuscript. This work was supported by the ‘Lendület’ programme of the Hungarian Academy of Sciences LP-2009-013/2012 (B.P.), LP-2012-32/2018 (C.P.) and LENDULET-BIOMAG (LP-2018-342, H.P.), the ELKH Lendület Programme LP-2017–10/2020 (C.P.), ELKH HAS-11015 (M.F.), the Wellcome Trust WT 098016/Z/11/Z (B.P.), The European Research Council H2020-ERC-2014-CoG 648364-Resistance Evolution (C.P.) and H2020-ERC-2019-PoC 862077–Aware (C.P.), H2020 ERAPERMED-COMPASS (DiscovAIR, H.P.), Chan Zuckerberg Initiative (Deep Visual Proteomics, H.P.), the National Research, Development and Innovation Office and the Ministry for Innovation and Technology under the “Frontline” Programme KKP KH125616 and 126506 (B.P. and C.P.), GINOP-2.3.2-15-2016-00026 (iChamber, B.P., P.H.), GINOP-2.3.2–15–2016–00014 (EVOMER, C.P. and B.P.), GINOP-2.3.2–15– 2016–00020 (MolMedEx TUMORDNS, C.P.), GINOP-2.3.2-15-2016-00006 (H.P.), GINOP-2.3.2-15-2016-00037 (H.P.), GINOP-2.3.2-15-2016-00044 (M.F.), National Laboratory of Biotechnology Grant NKFIH-871-3/2020 (B.K. and C.P.), and National Research, Development and Innovation Office grant FK-135245 (B.K.), The European Union’s Horizon 2020 research and innovation programme under grant agreement No 739593 (B.P and B.K.), János Bolyai Research Fellowship from the Hungarian Academy of Sciences (BO/352/20, B.K.), New National Excellence Program of the Ministry of Human Capacities (UNKP-20-5-SZTE-654, B.K.), and from the Chan Zuckerberg Initiative (Deep Visual Proteomics). B.Sz. holds a Premium Postdoctoral Fellowship of the Hungarian Academy of Sciences.

## Supplementary Figures

**Figure S1.**
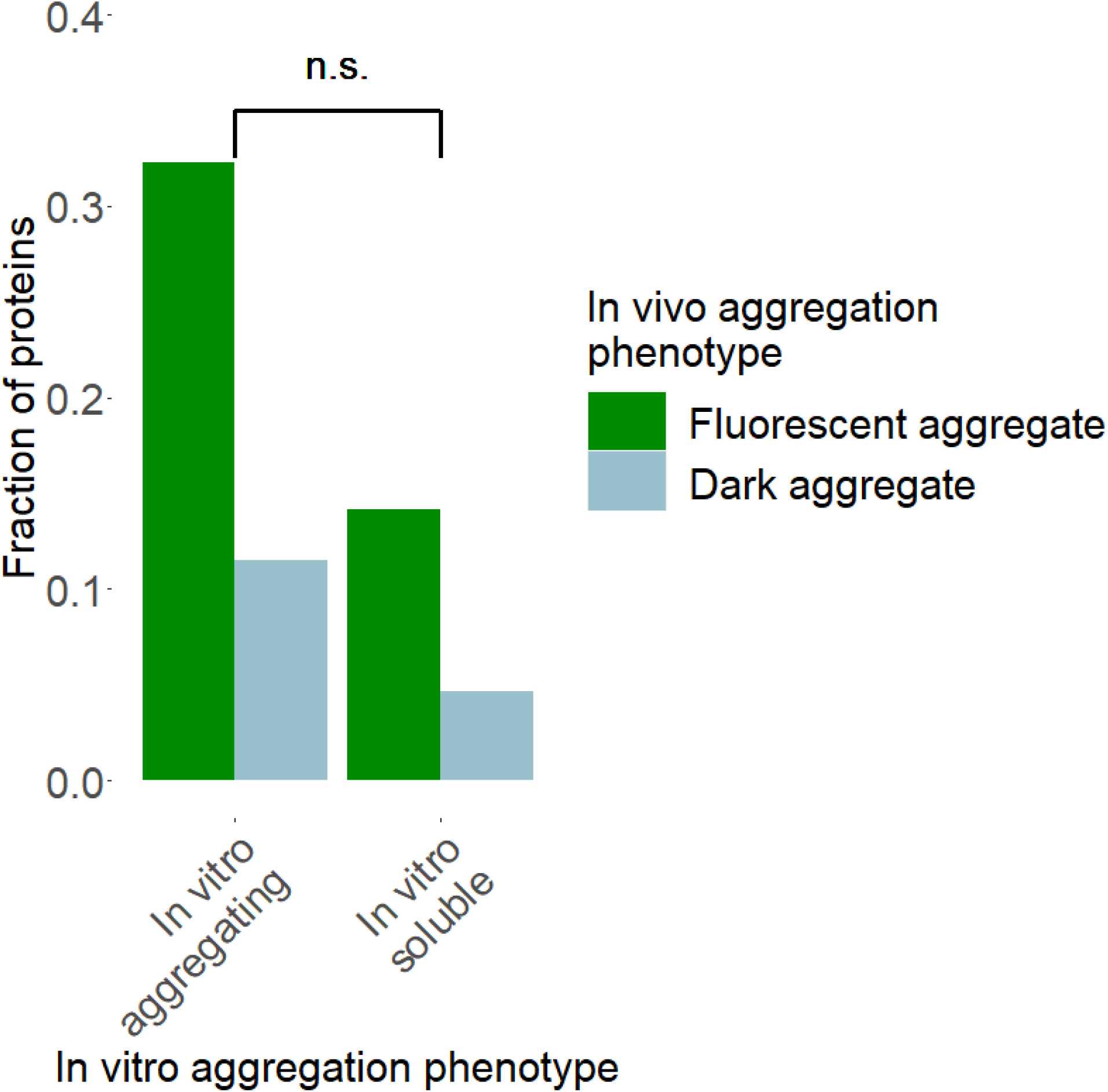
The two in vivo aggregation phenotypes show similar overlaps with in vitro aggregation. The relative frequencies of fluorescent versus dark aggregates are similar in the in vitro aggregating and in vitro soluble classes of proteins (P = 0.1873, Fisher’s exact test).

**Figure S2.**
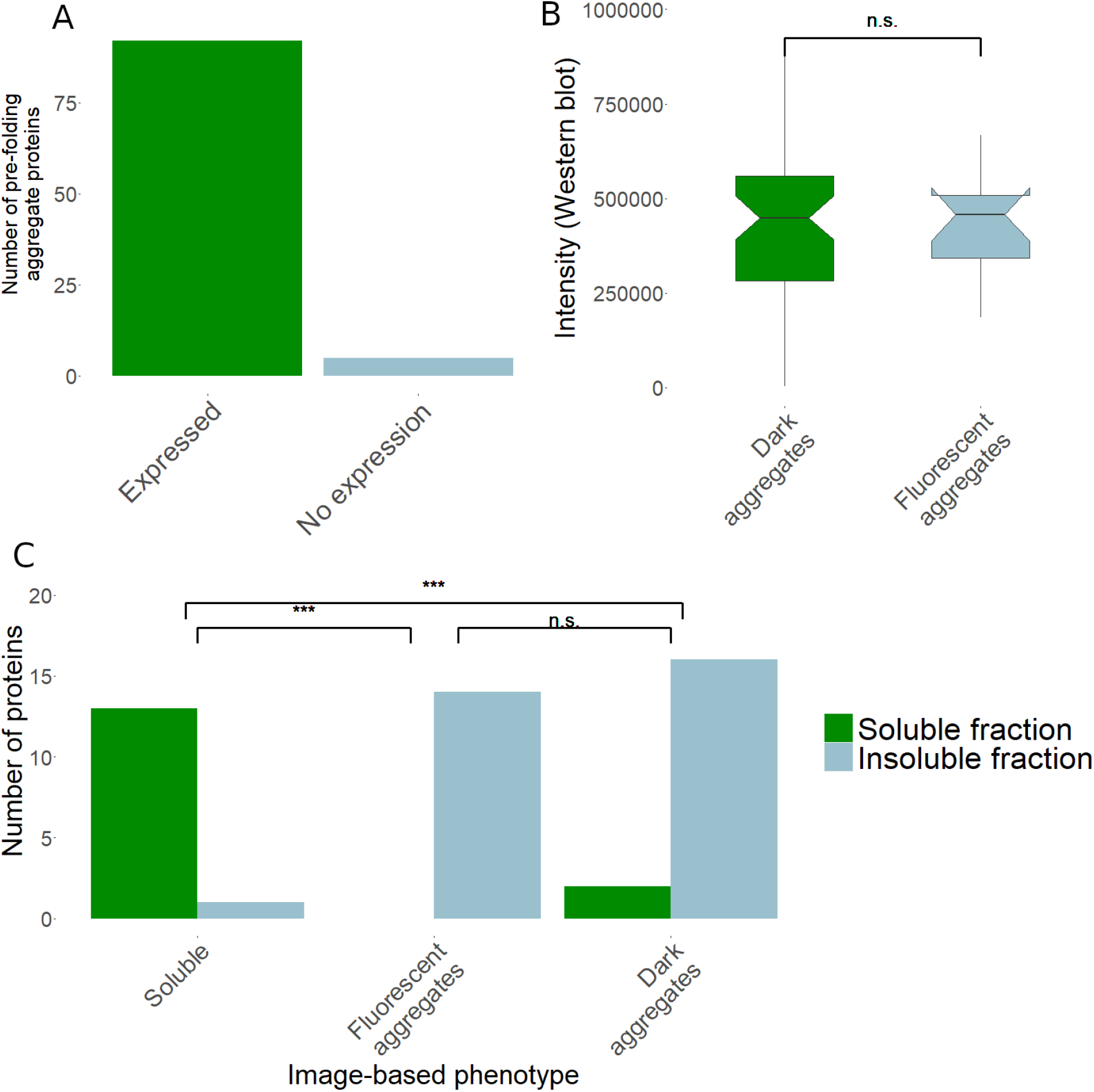
Experimental validation of protein overexpression and solubility phenotypes. **(A)** Summary of western blot analysis of a representative subset of 97 proteins classified as ‘dark aggregate’ in the image-based in vivo screen. 94.8% of these proteins are expressed as detected using western blot. **(B)** Quantified band intensity measurements from western-blot analysis of fluorescent and dark aggregate proteins. No significant difference was found between the band intensities (P = 0.8794, Wilcoxon Signed rank test, N = 71, see supplementary methods). **(C)** Validation of the high-throughput image-based cell classification by analyzing the soluble and insoluble protein fractions of a representative set of overexpression strains with SDS-PAGE and western blotting. The set of overexpression constructs represent 14, 14 and 18 proteins that show soluble, fluorescent aggregate and dark aggregate phenotypes, respectively, in the image-based screen. Proteins were classified as aggregate if no protein was detected in the soluble fraction using SDS-PAGE western blot (see Methods). Proteins showing solubility in the image-based analysis are significantly enriched in the soluble fraction compared to proteins that form fluorescent and dark aggregates (P = 7.48*10^−7^ and P = 5.26*10^−6^, respectively, Fisher’s exact test, N = 46). Proteins in fluorescent and dark aggregates show similar relative frequencies of protein aggregation (P = 0.49, Fisher’s exact test).

**Figure S3.**
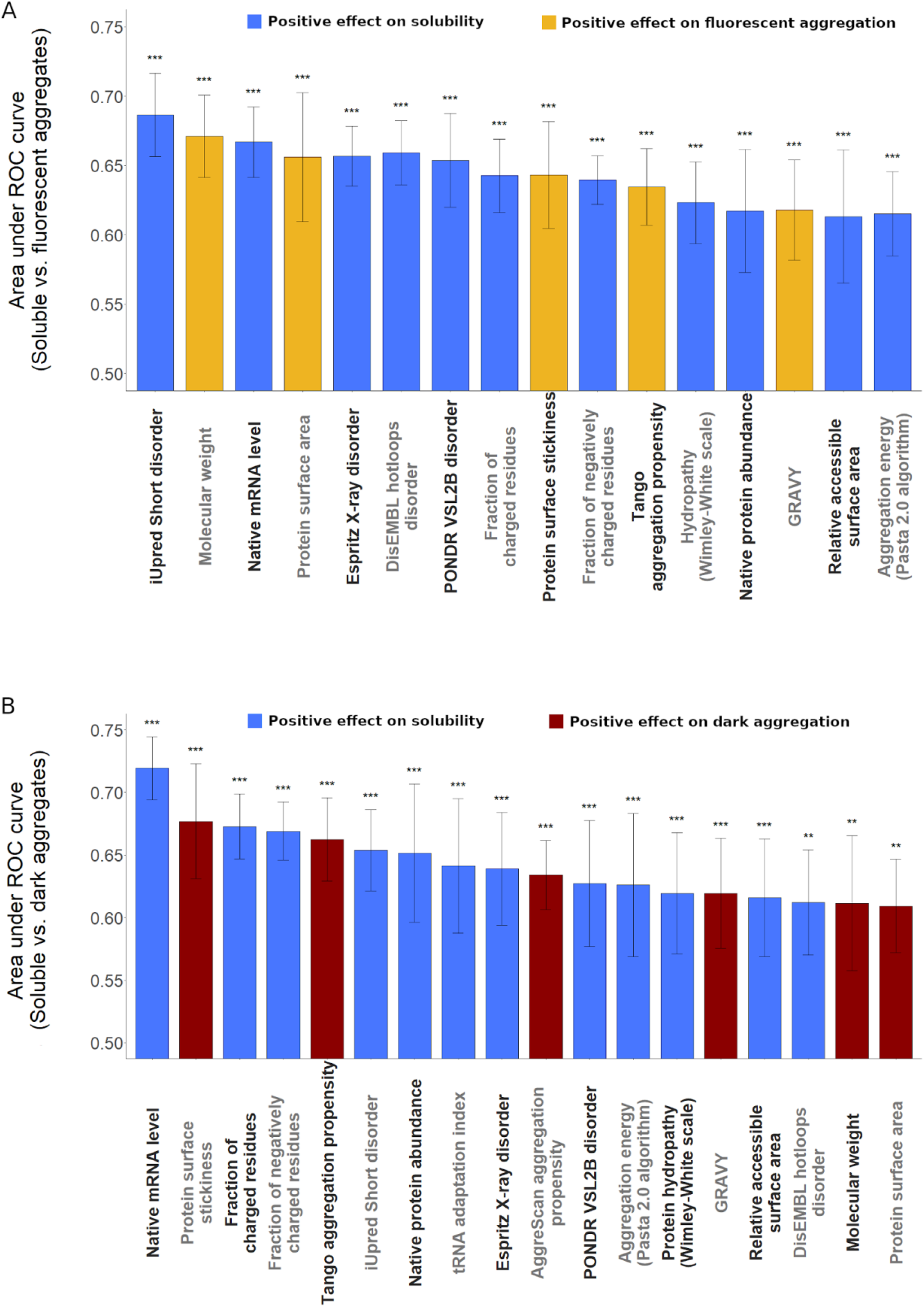
Major protein features discriminating soluble proteins from the two classes of aggregates. Features discriminating between proteins that remain soluble and those that form fluorescent (A) or dark (B) aggregates, respectively. The predictive ability of each feature was measured as the average area under the receiver operating characteristic (ROC) curve in a tenfold cross-validation procedure based on logistic regression analyses. All displayed protein features are statistically significantly predictive after adjustment for multiple testing using the false discovery rate method,*** corresponds to p adj <0.0001, ** corresponds to p adj < 0.01, * to p adj < 0.05 (logistic regression). Error bars show the 95% confidence interval for the AUC value of each feature.

